# Limited reciprocal surrogacy of bird and habitat diversity and inconsistencies in their representation in Romanian protected areas

**DOI:** 10.1101/2021.05.07.443068

**Authors:** Julia C. Geue, Paula J. Rotter, Caspar Gross, Zoltán Benkő, István Kovács, Ciprian Fântână, Judit Veres-Szászka, Cristi Domşa, Emanuel Baltag, Szilárd J. Daróczi, Gábor M. Bóné, Viorel D. Popescu, Henri A. Thomassen

**Affiliations:** Comparative Zoology, Institute for Evolution and Ecology, University of Tübingen, Tübingen, Germany; Institute of Medical Genetics and Applied Genomics, University of Tübingen, Tübingen, Germany; Romanian Ornithological Society / BirdLife Romania, Cluj-Napoca, Romania; Milvus Group, Bird and Nature Protection Association, Târgu Mureș, Romania; Department of Biological Sciences and Sustainability Studies Theme, Ohio University, Athens, Ohio, USA; Centre for Environmental Research, University of Bucharest, Bucharest, Romania; Marine Biological Station ”Prof. Dr. Ioan Borcea”, Agigea, “Alexandru Ioan Cuza” University of Iași, Iași, Romania

## Abstract

Because it is impossible to comprehensively characterize biodiversity at all levels of organization, conservation prioritization efforts need to rely on surrogates. As species distribution maps of relished groups as well as high-resolution remotely sensed data increasingly become available, both types of surrogates are commonly used. A good surrogate should represent as much of biodiversity as possible, but it often remains unclear to what extent this is the case. Here, we aimed to address this question by assessing how well bird species and habitat diversity represent one another. We conducted our study in Romania, a species-rich country with high landscape heterogeneity where bird species distribution data have only recently started to become available. First, we prioritized areas for conservation based on either 137 breeding bird species or 36 habitat classes, and then evaluated their reciprocal surrogacy performance. Second, we examined how well these features are represented in already existing protected areas. Finally, we identified target regions of high conservation value for the potential expansion of the current network of reserves (as planned under the new EU Biodiversity Strategy for 2030). We found that bird species were a better surrogate for habitat diversity than vice versa. Highly ranked areas based on habitat diversity were represented better than areas based on bird species, which varied considerably between species. Our results highlight that taxonomic and environmental (i.e., habitat types) data may perform rather poorly as reciprocal surrogates, and multiple sources of data are required for a full evaluation of protected areas expansion.

## Introduction

The ultimate goal of conservation prioritization is the protection of biodiversity at all levels of organization [1]. However, limited financial resources and competing stakeholder interests constrain the area that can reasonably be protected. The process of identifying potential regions for designation as protected area (PA) should therefore be undertaken thoroughly and strategically [2, 3, see 4 for a review]. The striking obstacle is however that biodiversity is very complex and difficult to characterize [5], and surveying biodiversity in its entirety is nearly impossible. Shortcuts necessarily need to be taken to quicken the prioritization process and ensure its feasibility [6]. One of these shortcuts is using a biodiversity or environmental indicator as a conservation surrogate [see 4 for a review, 7], which is: “An ecological process or element (e.g., species, ecosystem, or abiotic factor) that [should] […] represent (i.e., serve as a proxy for) another aspect of an ecological system” [8]. The efficacy and efficiency of surrogates for overall biodiversity (known and unknown) have progressively been evaluated [7, 9-13], and appear to be influenced by factors such as the size of the study area, type of surrogate, and the spatial resolution of surrogate data [e.g. 13, 14, 15]. Nevertheless, it often remains ambiguous to what extent a surrogate represents other levels of biodiversity, in particular across different levels of organization.

Biodiversity surrogates are usually subdivided into taxonomic and environmental surrogates [7, 10, 15]. Many studies have evaluated the efficacy of taxonomic surrogates for other taxonomic groups [e.g. see 6, 16 for a review, 17-21]. The general consensus is that one taxonomic group alone might not be an adequate surrogate for others [13, 14, 22, see 23 for a review, 24, 25], and the identification of PAs should include more than one species or taxonomic group [14, 26]. Yet again, for many areas in the world accurate species distribution data is scarce. One of the taxonomic groups for which rich datasets are available are birds, because they are of interest to many people and are therefore one of the best surveyed taxa in the world [26-28]. As such, birds are frequently used as biodiversity indicators and conservation surrogates, and their surrogacy effectiveness varies from representing overall species diversity well (other taxa than birds) [9, 23, 26, 29], or threatened birds being adequate surrogates for non-threatened bird species [14], to being poor surrogates for other taxa [20, 30, 31]. Adding more taxa [26] or even different biodiversity features, such as environmental diversity [13, 32], increased the overall surrogacy of birds for other levels of biodiversity.

Environmental diversity (ED), in particular habitat diversity, has the potential to be a powerful surrogate and represent other levels of biological organization, because habitat data can be generated quickly and relatively inexpensively from remotely sensed or extrapolated ground data [7, 15, 23, 33-36]. By using ED in conservation prioritization, it is assumed that selected areas do not only cover a wide range of different environmental conditions, but also areas, rich in other biodiversity features, such as species [35]. Furthermore, environmental surrogates may capture interactions between species and their environment [32], and compensate for a potential lack of congruence between taxonomic surrogates [22]. However, compared to taxonomic surrogates, the application of environmental surrogates received less attention. One potential reason may be that so far no clear and explicit recommendation about the efficiency of ED as a surrogate for biodiversity in general exists [e.g. 37, 38]. Moreover, there is no consensus on the way that ED is measured and included in a surrogacy approach, e.g. continuous versus discrete measures of ED [e.g. 37]. Some studies suggested that continuously distributed environmental variables (e.g. climate variables such as temperature and precipitation, or vegetation characteristics such as percent tree cover) may outperform discrete (classified) environmental data [e.g. 39, 40], but also may be inadequate [13, 23, 38] or at most better than random surrogates for species occurrence [7, 36]. Other studies, however, found that categorical environmental data in the form of pre-classified information (e.g. land classes, ecological vegetation classes or habitat types), may nevertheless be better surrogates than continuous environmental data [e.g.15, 32, 37, 38]. The representation of habitat or land cover categories for other levels of biodiversity may vary considerably, for instance being weak for plant species [32, 41], but better for plants than for vertebrates [10, 13, 15, 42, 43]. Yet, such contrasting results could also result from differences in the spatial extent and resolution of the study area, as well as the type of environmental data used and the amount of different environmental features included as a surrogate (vegetation or climate-based) [4, 15, 36, 44, 45].

Given uncertainties surrounding the potential for categorical habitat data to serve as a surrogate for biodiversity, the goal of our study was to evaluate its representation of one of the most frequently used biodiversity surrogates, bird species distributions, and vice versa. We implemented this analysis for Romania, a country within the European Union exhibiting high bird species and habitat diversity, likely caused by the variety of biogeographic regions it comprises [46, 47]. While 23% of Romania is protected, either under the pan-European Natura 2000 network or as natural or national parks or biosphere reserves [48], and despite its high levels of biodiversity, efforts to identify conservation priorities and evaluate the efficacy of the network of reserves to protect biodiversity have been sparse [mentioned by 11 but not examined, 21, 48-51]. One reason for this disparity is that species distribution data have only recently become widely available. As such, PA management could greatly benefit from prioritization efforts using systematic conservation planning principles and the latest available data, particularly when establishing new PAs [48, 49]. The implementation of such scientific research in the establishment and governance process of PAs is, however, often limited [52, 53]. This is not a unique situation, as for instance Natura 2000 sites consist of a diverse array of reserves designed for particular species, but not to protect biodiversity as a whole, so they often represent species and habitat diversity only to a limited extent [38, 49, 54, 55]. Furthermore, the European Commission decided to set new targets for 2030 and increase the percentage of protected areas in EU member states to 30% [56]. Hence, there is a need to identify additional areas for protection, which is best done using the principles of systematic conservation planning [57].

In this study, we first evaluated whether breeding bird species and habitat diversity based on remote-sensing data are adequate surrogates for one another. We assessed surrogacy of the two datasets using high-resolution data (1km) of (a) 137 modelled breeding bird species distributions and (b) 36 classes of mapped habitat types from the “Ecosystem Types of Europe” (ETE) data set [58]. Second, we evaluated whether existing protected areas (national and natural parks, biosphere reserves, wetland reserves and SPAs (as part of the Natura 2000 network)) in Romania are effective in representing areas of conservation concern for both birds and habitats. Finally, we identified additional areas that could be prioritized in an effort to expand the current PA network to more comprehensively protect bird and habitat diversity.

## Methods

### Study region

Romania is located in Eastern Europe, at the western shores of the Black Sea. It covers 238 397 km^2^ and natural landmarks and borders are dominated by the Carpathian Mountains and the Danube River. Five biogeographical regions have been characterized across Romania: Pannonian, Continental, Alpine, Steppic, and Black Sea. The heterogeneous landscape consists of an alternation between intensive and extensive agricultural areas and (semi-) natural areas, such as forest, open woodland, and grassland. As a member of the European Union, Romania is bound to the directives of the Natura 2000 network, and dedicated about 23% of its total landscape to conservation. The Natura 2000 network is an important biodiversity conservation measure [11], and consists of different types of protected areas: the terrestrial Special Protection Areas (SPA, for bird protection only), the terrestrial Sites of Community Importance, and Special Areas of Conservation (SCI and SAC, for habitats and/or species) [59]). In addition to, but partly overlapping with the Natura 2000 network, Romania also implemented protected areas designated as natural and national parks as well as biosphere reserves [48].

### Biodiversity features

#### Bird species distributions

Bird species occurrence data (from the years 2006-2018) were obtained from the forthcoming Romanian Breeding Bird Atlas ([60], in preparation), a scheme run by Milvus Group Association and the Romanian Ornithological Society. Because data based on species atlases often entail a geographic sampling bias [61], we first modelled the distributions of 137 breeding bird species using MaxEnt 3.4.1. [62] at two different resolutions (1×1km and 2×2km), depending on the species’ ecology or in some cases by the available data (S1 Table). In-depth details on the species distribution modeling approach are provided in S1 Supporting information.

#### Habitat types

We used the published maps of habitat types classified in the “Ecosystem Types of Europe” (ETE) data set (version 3.1)[58]. ETE is a combination of the non-spatially referenced EUNIS (European Nature Information System) habitat classification scheme and a spatially explicit habitat data set, the Corine-based “Mapping and Assessment of Ecosystem and their services (MAES)” ecosystem classes [63]. In Romania, 42 ETE habitat classes are mapped (level 2 classification) at 100 m resolution (S2 Table). Habitat classes including highly built-up areas (six classes) were excluded in the subsequent spatial conservation mapping. These built-up areas where selected according to the ETE classification category “J” (J1-J6, see S2 Table), which include buildings in cities and villages, industrial sites, transport networks, artificial water structures and waste deposits.

To produce maps of habitat types that match the spatial resolution of those for the bird species, we split the pre-defined ETE data set into single data layers per class (36 in total) and calculated the proportion of each habitat type within 1 km^2^ grid cells.

#### Data handling

All spatial data layers were re-projected to the Dealul Piscului 1970/ Stereo 70 projection and processed at a 1 km resolution containing a total number of 381 248 grid cells. Species distribution models at 2 km resolution were resampled to 1 km grid cell size. Preparation of input maps and post-processing of results was done in R (version 3.6.1), using the packages (zonator, raster, rgdal, rgeos, sp, maptools, tiff, data.table, plyr, dplry, ggplot2, zoo). Maps were visualized in QGIS (version 3.10.6 ‘A Coruña’).

### Spatial conservation prioritization

We prioritized areas for conservation using the software Zonation 4.0 [64]. Zonation can handle large data sets [65] and provides a priority ranking over the entire landscape rather than satisfying a specific target. The ranking is produced by iteratively discarding locations (grid cells) with the lowest conservation values, retaining the ones with the highest conservation value throughout the process [66, 67].

We used the additive-benefit function (ABF), which directly sums up the conservation value across features [68] and results in a reserve network with high average performance across all features [69]. The ABF algorithm is appropriate for our study since we aim to identify areas representing overall richness rather than core areas that lead to the equal representation of both common and rare species or habitats. The algorithm accounts for the total and remaining distributions of features, and optional feature-weights can be implemented [13]. We equally weighted habitat types and bird species distributions at the aggregate level to avoid prioritization biases due to the different numbers of features contained within (e.g., combined weights for 137 bird species or for 36 habitat types summed to 1). To exclude land uses that for administrative or ecological reasons did not contribute to either overall conservation value or to the expansion of protected areas (six classes of built-up area), we applied a cumulative negative weight of -1 to these layers [68].

Performance curves were produced with the R package ‘zonator’ [70]. These curves show the proportion of the original occurrence of features remaining in the landscape as a function of the proportion of the landscape that is lost [71]. The curves start at 1.0, where the entire distribution of features is represented in the full landscape, and end at 0.0, where the entire landscape is lost.

Because we observed a wide spread in the performance curves of the bird species, we explored potential underlying patterns related to their broad habitat requirements. We grouped species into their preferred breeding habitat types (S1 Table) to assess differences between groups and their performance when the prioritization is accounting for all bird species.

We also suspected that the range size of feature types, in particular within bird species, influences their performance in the prioritization. Specifically, we assumed that range restricted species would perform better, since these species might be retained throughout many prioritization iterations. Yet, this may only be the case when range-restricted features largely overlap with more widespread features. To explore this further, we calculated the AUC (area under the curve) of each feature performance curve, and plotted these as a function of range size (Fig 1). For bird species we calculated range sizes by summing the Maxent probabilities, and for habitat types we summed the area in km^2^.

**Figure 1.**
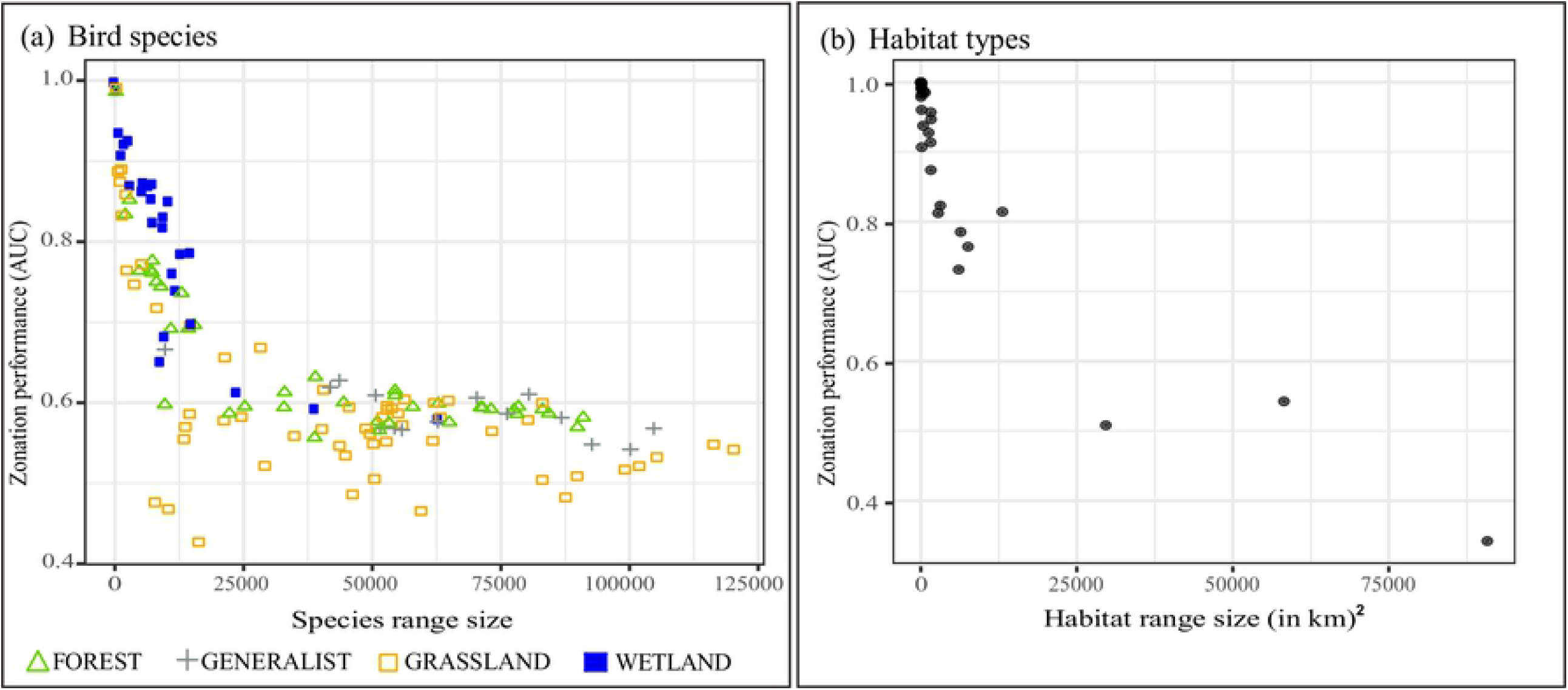
The Zonation performance of individual features (AUC) as a function of its corresponding range size. (a) Individuals bird species, belonging to one of the four breeding habitat groups. Green triangle = forest to (dense) woodland; grey cross = generalist and close to humans; yellow square = arable land, open woodland to grassland; blue square = wetland and shores. The values for the range sizes of bird species were computed by adding up Maxent species distribution values. (b) Individual habitat types

### Surrogacy analyses

We evaluated the reciprocal surrogacy of bird species and habitat types, and assessed the efficacy of the existing network of protected areas to protect these biodiversity features. To test the surrogacy of the two feature types, we ran separate analyses using one feature type as the surrogate and the other as the target. To do so, bird species and habitat types were both included in each run, but positive weights (=1) were only assigned to the surrogate, while the target was assigned a weight of 0.

We evaluated the surrogacy power of each feature type using the performance curves. A performance curve by itself provides, however, little information, and for correct interpretation it should be compared to an optimal and a random curve [23]. For instance, when testing whether habitat types are a good surrogate for bird species, the optimal curve is equivalent to the surrogacy of bird species for themselves. The random curve in this scenario reflects the representation of bird species expected in the absence of biological data, when ‘area’ is used as a surrogate [72]. Qualitatively the surrogacy value can be assessed visually by comparing the three curves. The closer the target curve is to the optimal curve, the higher the surrogacy value. To quantify the surrogacy power, we calculated an equivalent to the species accumulation index (SAI; Ferrier (73)):

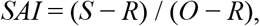

where *S* is the area under the target curve, *R* is the area under the random curve, and *O* is the area under the optimal curve. The optimal curve was extracted from the runs when targets were used as a surrogate themselves. To create the random curve, we executed 100 surrogacy runs with randomly, uniformly distributed data as a surrogate and bird species and habitat types as targets. We used the mean of the corresponding target curves to calculate SAI.

### Evaluation and potential expansion of the protected area network

To evaluate the representation of habitat and birds in existing reserves, we specifically focussed on SPAs, national and natural parks, and biosphere reserves. We thus excluded the SCI and SAC areas (Natura 2000 sites), since they are designed to protect specific species or habitats, but do not necessarily protect others - or even biodiversity as a whole. To evaluate the effectiveness of the current network in Romania, we tested 1) how well current PAs represent areas of conservation concern for bird species and habitat types, and 2) how much of the individual feature type’s distributions are represented within the current network. Furthermore, we 3) assessed which areas should be prioritized when expanding the current conservation network.

The analyses for 1) and 3) were based on a Zonation prioritization outputs, where both bird species and habitat types had been considered simultaneously. We did not differentiate between protection levels of the existing PAs. If PAs had been selected indiscriminately, we expected that Zonation values within PAs would be uniformly distributed, as they are across the entire study region. We thus tested the frequency of Zonation values within PAs against a uniform distribution using a Chi-square test. For 2) we summarized the distribution of bird species and habitat types within current PAs as a proportion of their total distribution via boxplots (S1 Fig b and c).

To identify potential areas that should be prioritized when expanding the current network of PAs, we performed a mask analysis [74]. In this analysis, current PAs are included as a mask layer, and are assigned a high rank (=1) in the final prioritization map. As such, the next highly ranked areas outside protected areas can be identified as potential expansion areas that represent bird and habitat diversity well.

## Results

### Spatial conservation prioritization

Both the separate and combined prioritization using bird species and habitat types resulted in broadly similar patterns, with highly ranked areas in the Carpathian Mountains, river valleys and parts of the Danube Delta. However smaller-scale differences are apparent, in particular with respect to the size and clustering of those areas (Fig 2a and S2 Fig).

**Figure 2.**
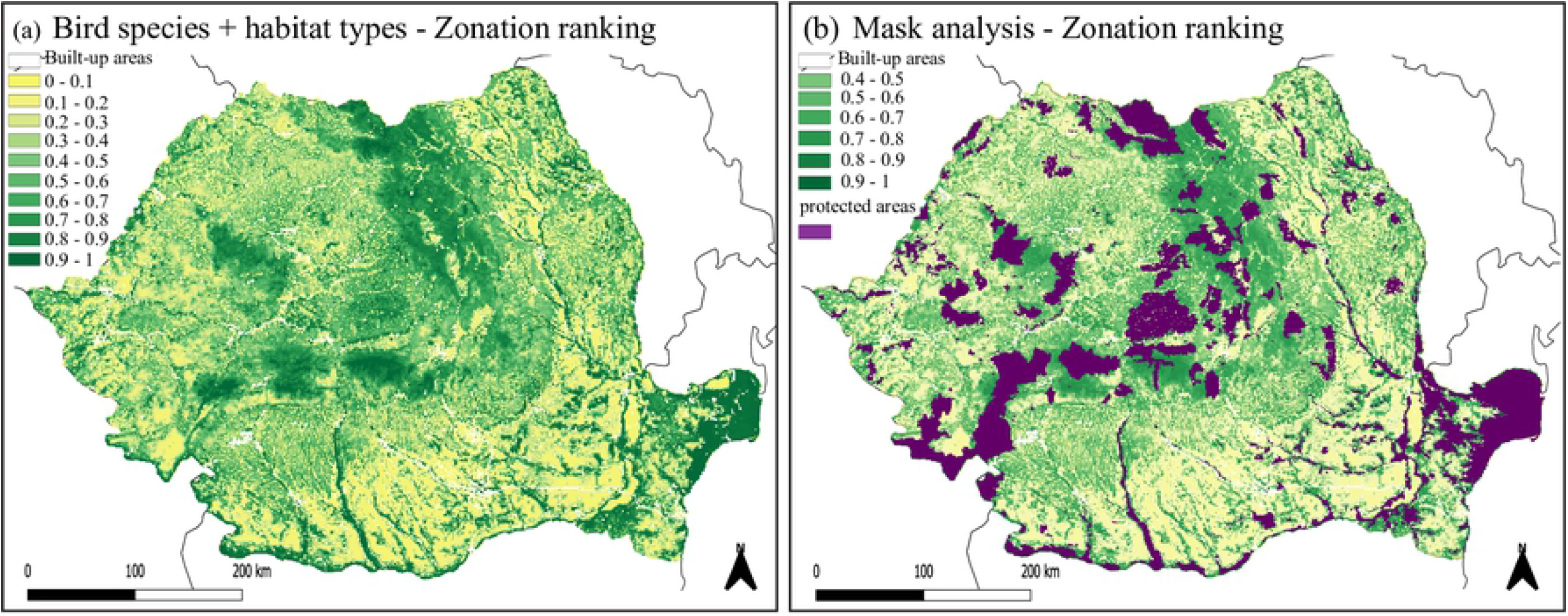
Study region with Zonation ranking based on bird species and habitat type data without (a) and with (b) considering currently protected areas (mask analysis). Colors indicate importance ranking scores for conservation, with 0 meaning lowest importance and 1 meaning highest importance. Built-up areas are indicated in white and were excluded from prioritization. Purple in panel (b) indicates current protected areas

The overall performance of bird species for themselves was rather low (AUC=0.65, area under the bird performance curve) (Fig 3a and S3 Table), but we observed considerable differences between groups based on breeding habitat (S3 Table).

**Figure 3.**
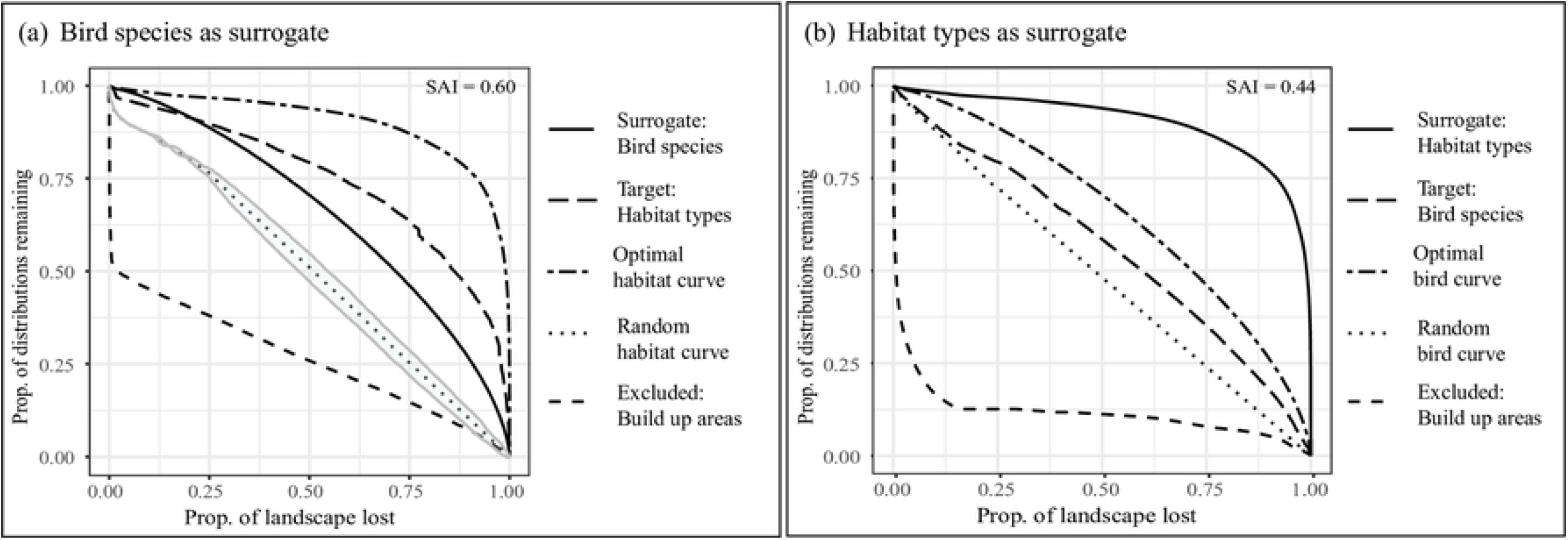
Performance and surrogacy curves quantifying the average proportion of original feature distributions represented as landscape is lost. Built-up areas were negatively weighted and hence excluded from the prioritization (lower dashed line). The area between the target curve and the random curve divided by the area between the optimal curve and random curve represents the efficacy of the surrogate (SAI; Species accumulation index). In panel (a) bird species were used as a surrogate for habitat types and in (b) habitats were used as a surrogate for birds

Wetland and shore-breeders were best retained through the ranking process, followed by those breeding in “forest to (dense) woodland” areas (Fig 4). In contrast, birds breeding in “arable land, open woodland to grassland” or being “generalist and close to humans” were lost much more quickly (Fig 4).

**Figure 4.**
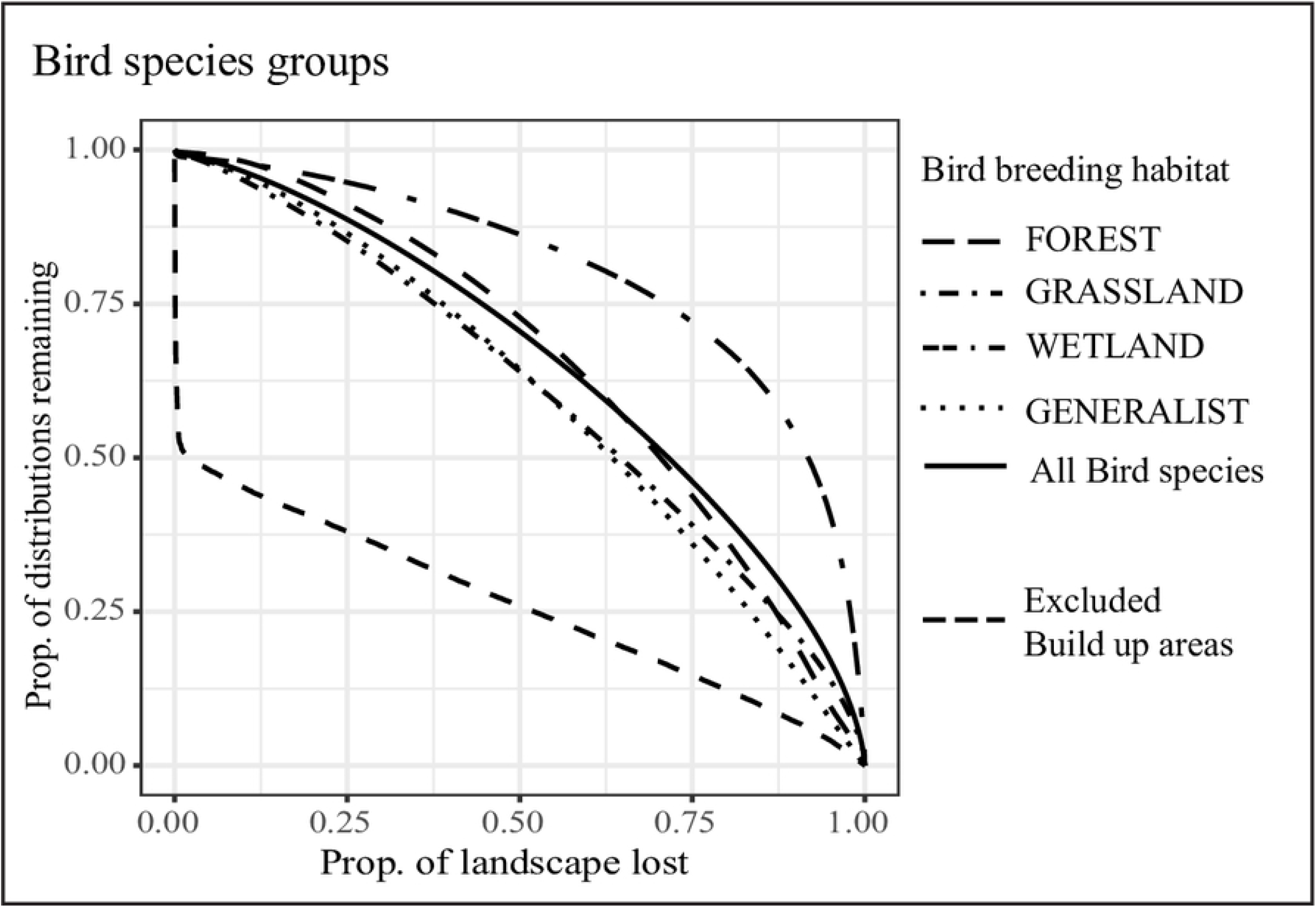
Performance curves for bird species split by breeding habitats. The solid line is the average performance curve of all bird species used in the surrogacy approach. Built-up areas were negatively weighted and hence excluded from the prioritization (lower dashed line)

To explore this further, we plotted each species’ performance as a function of its range size (Fig 1 and S1 Table), and found a clear negative trend. “Wetland and shore” breeders include more range-restricted species compared to other groups and at the same time performed best in the prioritization, whereas forest, generalist and grassland birds overall have larger ranges, and performed worst in the prioritization. In addition, the distributions of wetland and shore breeders often overlap with those of other groups, those resulting in areas of high species richness that are preferentially prioritized by the ABF algorithm (S3 Fig).

Habitat types were generally retained well throughout the prioritization process (AUC=0.9, area under the habitat surrogate curve) (Fig 3b and S3 Table). We observed that features with smaller ranges were retained the longest (Fig 1 and S2 Table).

### Surrogacy analyses

Birds were a moderately good surrogate for habitats (SAI = 0.60). Interestingly, birds represented habitats better than themselves (Fig 3a), although as shown above this is only true for the representation of all birds combined, and there are large differences between bird groups (Fig 4). The representation of habitats for birds on the other hand, was less effective (SAI = 0.44; Fig 3b).

### Evaluation of protected areas and identification of expansion regions

We found that the Zonation values within current PAs, when both habitat types and bird species were considered, differ significantly from a uniform distribution, with an overrepresentation of higher values (Chi^2^ test, Chi^2^ = 29289, df = 9, p-value < 2.2e-16) (S1 Fig a). These results suggest that current PAs generally comprise areas of high conservation value better than would be expected based on a random assignment of areas for conservation. However, current PAs also comprise a considerable amount of land surface area with relatively low conservation values based on bird and habitat diversity, suggesting that improvements could be made.

Habitat types are relatively well represented in the current protected areas network (S1 Fig c), with the exception of grassland, heathland and woodland habitats. Among the breeding groups, generalist and grassland breeders are on average represented less well than expected under a random assignment, although in the grassland breeding group much variation between the species can be observed (S1 Fig b).

The mask analysis highlighted transition areas from highland to lowland regions, such as along the northern Carpathian Mountains, the eastern foothills of the Carpathian Mountains, and the eastern part of the Apuseni Mountains (Fig 2b) as particularly important expansion sites for bird and habitat conservation.

## Discussion

The necessity to rely on surrogates for conservation prioritization raises the question of how effective they are. Here we evaluated the mutual surrogacy power of bird species and habitat types in Romania, an area in Europe with high biodiversity, and demonstrated that neither birds nor habitat types are effective surrogates of one another. Birds represented 60% of habitat conservation priorities, while habitats were less effective at representing bird conservation priorities (44%). These results are not only concordant with studies in other regions suggesting to use more than one type of surrogate for conservation prioritization [13, 32, 42], but also point out that environmental data as conservation surrogates for species should be carefully evaluated prior to applications of protected area expansions. We also found that existing protected areas in Romania capture areas of high conservation value for both biodiversity features better than expected at random, but could potentially be designed more effectively and more efficiently. Finally, we identified additional areas that should be prioritized in case the existing network were to be expanded under the European Union Biodiversity Strategy to 2030, or where conservation strategies for conserving avian and habitat diversity on private lands could be incentivized.

### Bird species as a surrogate

The effectiveness of 137 breeding bird species as a surrogate for habitats was ∼60% of that of habitats for themselves (Fig 3a). Thus, in the absence of other data, birds could represent habitat types better than random, but only to a limited extent. These results appear robust because we included many bird species, breeding in a wide variety of habitats (S1 Table), thus covering the existing habitat diversity quite well. Our results corroborate other studies that found that taxonomic groups are poor surrogates for one another [for other taxonomic groups, e.g. 13, 14, 25, 26], and should be used cautiously as surrogates for habitat diversity.

Interestingly, when prioritizing bird species only (Fig 4) we found that wetland and shore birds were much better represented than forest, grassland, and generalist species. This unexpected result corroborates the focal areas of the Bird Directive, which demands particular attention to wetland species [75, Art 4 (2)]. A potential explanation for this representation bias is the emphasis of the additive-benefit function (ABF) on high average performance across all features - in the case of bird species, areas with high species richness [69] – combined with differences in range sizes between the bird groups [14, 67]. We found that species richness was highest in areas where the distributions of wetland-breeding species overlapped with those of species breeding in other types of habitat (S3 Fig). Because wetland birds generally have small ranges due to the limited availability of suitable habitat [76], Zonation prioritized the species-rich wetlands over areas with fewer species, where more widely distributed species occur (Fig 1). These results are in line with similar patterns in small versus large-range moths [20], butterflies, reptiles, and amphibians [14]. The representation bias in our study may be exacerbated by associations of generalist species to human-dominated landscapes. Because we negatively weighted and hence excluded built-up areas from the prioritization, species occurring in those areas may be underrepresented in the final results.

### Habitats as a surrogate

Habitats as a surrogate for birds were only 44 % as effective as the maximum possible. This result is consistent with other studies showing that environmental diversity may have limited suitability as a proxy for the diversity of small vertebrates (including bird species) [32, 77]. Yet, habitats represented birds better than random (Fig 3b), potentially due to the influence of habitat structure on bird species occurrence and distributions [43]. It remains unclear whether higher spatial and thematic resolutions – in particular more detailed habitat classifications – could improve the mutual representation.

The underlying concept of using environmental diversity (ED) as a conservation surrogate is that by selecting areas that cover a wide range of environmental conditions, other levels of biodiversity should be covered equally well [35]. The surrogacy power of ED may, however, depend on how it is tested and implemented in prioritization schemes, and whether it covers multiple taxonomic groups. Some studies suggested that pre-classified environmental data such as the ETE dataset [58] perform equally well or better than continuous environmental variables as a surrogate for species diversity [e.g. 15, 32, 37, 42, 78]. However, inconsistencies in the application of ED as a conservation surrogate and in what form it should be implemented (e.g. as discrete classes or continuous variation), are still unresolved, and also likely depend on the types of species data available. For instance, implementing ED as a distance matrix across as many environmental variables as possible, in combination with a different optimization procedure may in some cases help improve its surrogacy power [e.g. 36]. Yet, continuous environmental data such as climate or vegetation characteristics such as density or cover may fail to capture potentially important attributes summarized in classified habitat data like the ETE. To this end, more empirical evidence is needed from direct comparisons between methods as well as the performance of different measures of ED to provide a solid basis for recommendations of best practice.

Given the still existing uncertainties surrounding the use of ED, we set out with pre-classified habitat types, which are themselves based on a wide range of climatic and environmental conditions [58]. Within the methodological and geographical scope of our study, results suggest that habitat classes performed relatively poorly at representing bird biodiversity in Romania, and ideally should not be used on their own in prioritization efforts. Instead, combining taxonomic and environmental surrogates could increase the surrogacy power for the protection of overall biodiversity [13, 42], but a single taxonomic group may not suffice. For instance, habitats and birds did not perform well in representing amphibians and reptiles in other areas [15, 38, 43]. Thus, we recommend to combine environmental and taxonomic surrogates, preferentially from multiple taxonomic groups.

### Representation in existing protected areas and conservation implications

We found that a considerable fraction of PAs is located in areas with high conservation values. It is important to stress, however, that our evaluations by no means suggest that the current network of PAs is sufficient. Around 23% of Romania’s land surface area is currently under protection, and improvements to the protected area network may be necessary [48, 49]. Large ecoregions and several widespread bird and mammal species may be protected sufficiently well, but smaller ecoregions, as well as invertebrate and plant species are for example underrepresented in the existing Natura 2000 network [49]. The current network of PAs consists of reserves designed for various purposes. In our evaluation, we specifically focused on those that have been designed to protect birds, habitats, or biodiversity as a whole, i.e. SPAs, national and natural parks, and biosphere reserves. We found that these PAs represent areas of high bird or habitat conservation value better than a random assignment of areas for protection. However, habitats were better represented than birds (S1 Fig b and c). We also found that rare habitats are well represented, which is consistent with results for the Czech Republic [54]. These habitats typically are wetlands and shores, large areas of which are protected in the Danube Delta. Surprisingly, the representation of grassland and woodland habitats was rather poor. A likely reason for this result is the large area of wood- and grassland habitats in Romania, only part of which can be represented in PAs (S4 Fig). In contrast, rare habitats such as littoral areas are represented at high percentages, because they can be entirely contained within a fraction of the total land surface area. Despite the fact that current PAs capture important areas for conservation relatively well, a tail of areas with low conservation value can also be observed (S1 Fig). It remains unclear whether these areas may be important for other reasons, such as for other taxonomic groups, or as corridors between areas of high conservation value. Yet, the presence of areas with low conservation value also suggests that improvements in both the efficiency and efficacy of the network may be possible. To this end, we identified areas that should be prioritized based on bird and habitat diversity in a scenario of future expansions of the current network. A recent study suggests that such improvements may best be developed at the level of Biogeographical Regions rather than at the national level [57].

Protected areas are a crucial component of conservation, but the identification and designation of PAs is often a lengthy and difficult process. In addition, even when the new targets for the EU Biodiversity Strategy are met, 70% of the land surface area will remain unprotected. Hence, effective conservation also depends on the protection of biodiversity outside of PAs. To do so, the development of incentives for targeted management practices to retain high diversity of species and habitats should be prioritized [79], yet scientific research that can support management decision is largely lacking [80].

We conducted this study in Romania, a heterogeneous and species-rich country, comprising five biogeographical regions. We exploited an extensive data set on breeding bird species that has only recently become available for the country. Our study adds to the body of evidence that taxonomic and environmental surrogates represent one another only to a limited extent. Hence, the use of just one type of surrogate may not capture the broad patterns of biodiversity sufficiently well. This situation is less than ideal, as conservation measures respond to the biodiversity crisis, with little time to collect data on the distribution of species or habitats. Although these data are becoming increasingly available, our results highlight the need for investing in survey and monitoring schemes in countries such as Romania, where data still remains relatively scarce. Our study also presents an example of the importance of scientific research in informing conservation strategies as a stakeholder, which often remains underrated [52, 53].

## Supporting information - Figures captions

**S1 Figure** (a) Barplot of conservation values of areas in current reserves. The horizontal dashed line indicates the expected frequency of each conservation value (freq = 3338.6), had the current PAs be selected at random. The high frequencies of high conservation values, combined with the low frequencies of low conservation values suggest that current PAs were selected efficiently. (b-c) Box-and-whisker plots for birds (b) and habitats (c) showing the proportion of the total distribution of each group of feature types that is represented in the existing protected area network. A dotted line indicates the random expectation for the representation of each feature class based on the amount of protected area in Romania (∼ 23% of land surface area)

**S2 Figure** Study region with Zonation ranking based on (a) Bird species and (b) habitat types. Colors indicate importance ranking scores for conservation, with 0 meaning lowest importance and 1 meaning highest importance. Built-up areas are indicated in white and were excluded from prioritization

**S3 Figure** Overlapping bird species occurrences per breeding habitat group: (a) forests to (dense) woodland, (b) generalist and close to humans, (c) arable land, open woodland to grasslands, and (d) wetlands and shores. Red indicates species-rich areas; white to grey indicate no or low overlap of species occurrences

**S4 Figure** Study region with (a) forest habitats and (b) grassland habitats highlighted. The used protected area network is highlighted in grey

